# Viral protease NIa-Pro increases MEDIATOR SUBUNIT16 cleavage and plant susceptibility to turnip mosaic virus and its aphid vector *Myzus persicae*

**DOI:** 10.1101/2023.10.24.563845

**Authors:** Swayamjit Ray, Tyseen Murad, Gabriella D. Arena, Kanza Arshad, Zebulun Arendsee, Venura Herath, Steven A. Whitham, Clare L. Casteel

## Abstract

Plant viruses, such turnip mosaic virus (TuMV), both trigger and inhibit host plant defense responses, including defenses that target their insect vectors, such as aphids. TuMV infection and its protein, NIa-Pro (nuclear inclusion protease a), suppress aphid-induced plant defenses, however the mechanisms of this suppression are still largely unknown. In this study, we determined that NIa-Pro’s protease activity is required to increase aphid performance on host plants and that 40 transcripts with predicted NIa-Pro cleavage sequences are regulated in Arabidopsis plants challenged with aphids and/or virus compared to healthy controls. One of the candidates, MEDIATOR 16 (MED16), regulates the transcription of ethylene (ET)/jasmonic acid (JA)-dependent defense responses against necrotrophic pathogens. Using immunoblots, mutants, bioassays, and qRT-PCR, we show that a nuclear localization signal is cleaved from MED16 in virus-infected plants and in the presence of NIa-Pro along with the presence of aphids, suggesting MED16 functions in the nucleus may be impacted. Consistent with this, aphid induction of the MED16-dependent transcript of *PLANT DEFENSIN 1.2 (PDF1.2)*, was reduced in virus-infected plants and in plants expressing NIa-Pro compared to controls, and NIa-Pro’s protease activity was required for this reduction. Finally, we show performance of both the virus and the aphid vector was enhanced on *med16* mutant Arabidopsis compared to controls. Overall, this study demonstrates MED16 regulates defense responses against both the virus and the aphid and provides insights into the mechanism by which TuMV suppresses anti-virus and anti-herbivore defenses.

## Introduction

Proteins and small molecules that are used by pathogens and insect herbivores to successfully colonize a plant are known as effectors, and these molecules can also serve as reliable cues of attack that are recognized by the plant (Jones and Dangl, 2006; Acevedo *et al*., 2015; Ray and Casteel, 2022). While plant recognition of effectors, effector-mediated suppression of plant defense, and plant counter defense have been extensively studied in recent years (Acevedo *et al*., 2015; Zhou and Zhang, 2020; Ngou *et al*., 2022), the mechanistic details of multi-trophic and multi-player effector interactions are still not fully understood (Ray and Casteel, 2022). For example, plants are confronted with effectors from viruses and insect vectors during vector-borne virus transmission. In some cases, plant defense genes are activated by insect feeding exclusively, but trigger defenses that target both the viral pathogen and the insect vector, such as with the leucine-rich repeat receptor (NLR) gene *VAT-1* in melon (Dogimont *et al*., 2014). There are also viral and insect effectors that can suppress plant defenses to increase the performance of a virus and the insect vector, such as with NIa-Pro (nuclear inclusion protease a) from turnip mosaic virus (TuMV) (Casteel *et al*., 2014) and the whitefly salivary protein Bsp9 (Nalam *et al*., 2019; Wang *et al*., 2019). On the other hand, some viral effectors antagonize vector performance while also benefiting the virus, such as 2b of cucumber mosaic virus (CMV) (Westwood *et al*., 2013), further complicating the dialog of attack, recognition, and response.

The processes of protein metabolism and turnover play pivotal roles in plant defense and are often targeted by insects and pathogens (Huang *et al*., 2012; Zhou *et al*., 2015; Bera *et al*., 2022). For example, in response to TuMV infection, Arabidopsis plants downregulate plant protease gene expression, while upregulating expression of genes involved in autophagy and protein turnover (Bera et. al, 2022). Pathogens also counter plant defenses by interfering with protein turnover, often through interactions with the ubiquitin proteasome complex (Banfield, 2015) (Üstün *et al*., 2013; Üstün *et al*., 2014). Furthermore, when insects and pathogens induce jasmonic acid signaling, several negative regulators of the pathway are targeted for degradation by the ubiquitin proteasome pathway (Katsir *et al*., 2008). This degradation process is thwarted by several viral effector proteins, such as C2 from tomato yellow leaf curl virus (TYLCV) and 2b from CMV, which stabilize the JAZ negative regulators, thereby blocking the downstream induction of plant defense (P., Li *et al*., 2019; Westwood *et al*., 2013). Viral effectors are also known to interact with the ubiquitin complex to mediate the suppression of host RNA silencing and for targeting plant-defense related proteins for degradation (Verchot, 2016).

Potyviruses encode a single polyprotein that must undergo precise cleavage events by viral encoded proteases to coordinate different aspects of viral replication and proliferation in the host cell (Carrington *et al*., 1993; Maia *et al*., 1996). Recently, it was shown that these viral proteases can also cleave plant proteins (Huogen *et al*., 2022), a well-known phenomenon in animal-virus systems (Zaragoza *et al*., 2006; H., Li *et al*., 2019; Visser *et al*., 2020). For instance, in *Nicotiana benthamiana* PHYTOCHROME-INTERACTING PROTEIN (AtPIF), a histone reader protein (AtEML2), and a lysine ketoglutarate reductase (AtDUF707) are cleaved by NIa-Pro (nuclear inclusion a protease) from plum pox virus (PPV), TuMV, and tobacco etch virus (TEV), all members of the *Potyviridae* family (Huogen *et al*., 2022). However, the specific function of these cleavages in plant-virus interactions was not determined. Additionally, it is estimated that NIa-Pro from TEV can interact with 76 different plant proteins (Martínez *et al*., 2016), suggesting many unknown mechanisms still need to be identified.

TuMV infection, and expression of its NIa-Pro, suppress aphid-induced defenses downstream of the ethylene signaling in plants (Casteel *et al*., 2014; Casteel *et al*., 2015; Bak *et al*., 2017), but the mechanisms of this suppression are still largely unknown. In this study, we hypothesized that NIa-Pro may suppress plant defenses by cleaving plant proteins that are required for aphid defense induction. We used site-directed mutagenesis to abolish the protease activity of NIa-Pro, which prevented NIa-Pro from increasing aphid fecundity on host plants. To identify potential plant proteins that could be cleaved by NIa-Pro, we screened the transcriptome of virus-infected Arabidopsis plants with and without aphids and found 40 candidate proteins, including MEDIATOR SUBUNIT16 (MED16). MED16 is known to regulate ethylene (ET)/jasmonic acid (JA)-dependent defense response in Arabidopsis (Wang et al., 2015). Using transient expression, stable transgenics, and immunoblots we demonstrated that in the presence of NIa-Pro or TuMV, MED16 abundance and it’s cleavage increases, while aphid induction of the MED16-dependent defense transcript *PLANT DEFENSIN 1.2* (*PDF1.2*) is suppressed (Zarei *et al*., 2011). Cleavage occurred largely in the cytoplasm and depended on the protease activity of NIa-Pro, suggesting NIa-Pro cleavage disrupts MED16-dependent transcription by blocking the nuclear localization of MED16. In support of this, we found that both the virus and the aphid vector performed better on *med16* mutant Arabidopsis compared to control plants, indicating that MED16 plays a crucial role in the defense response against aphids and virus.

## Materials and Methods

### Plants, insects, and virus infection

*Arabidopsis thaliana* (Col-0) and *Nicotiana benthamiana* plants were grown in Cornell mix [by weight 56% peat moss, 35% vermiculite, 4% lime, 4% Osmocote slow-release fertilizer (Scotts, Marysville, OH, USA, http://www.scotts.com)] in 16:8 light:day cycle in growth chambers at 24C. Arabidopsis mutant line *med16* was obtained from Arabidopsis Biological Research Center (ABRC) and the seeds of MED16-FLAG complementation line in *med16* mutant background was generously provided by Dr. Zhonglin Mou at University of Florida (Wang *et al*., 2015). *Myzus persicae* colonies were maintained on *Nicotiana tabacum* plants grown in the same growth conditions as stated above.

For virus infection, four-week-old Arabidopsis plants were rub-inoculated using stock plants of *N. benthamiana* infected with an infectious clone of TuMV containing the green fluorescent protein sequence (TuMV-GFP) (Lellis *et al*., 2002). Inoculum was prepared by grinding one gram of fresh leaf tissue from *N. benthamiana* in 5 ml of 0.02M potassium phosphate buffer (pH=5.2) on ice. The homogenized slurry was gently rubbed onto two rosette leaves of Arabidopsis plants with carborundum. As a control, inoculum was prepared from healthy *N. benthamiana* leaves, and used to inoculate a second set of plants at the same time (mock-inoculated). *Agrobacterium tumefaciens* bacteria expressing TuMV-GFP was cultured overnight in Luria Bertani media and cells were resuspended in 0.1M magnesium sulfate and 150µM acetyl syringone to obtain a final OD_600_ of 0.1. One milliliter of resuspended bacterial cells was then infiltrated into the ventral side of *N. benthamiana* leaves using a needle-less syringe. Plants that showed symptoms for viral infection and GFP expression after a week of infiltration were used as stock plants to rub-inoculate Arabidopsis plants.

### Cloning of constructs

The coding region of NIa-Pro protein was cloned from TuMV into the binary vector pMDC32 downstream of the 35S promoter as well in pSITE vector using Gibson assembly (New England Biolabs, USA)(Gibson *et al*., 2009). Protease activity of NIa-Pro was abolished by mutating cysteine 151 to alanine (C151A) using the same construct and the QuikChange Lightning Site-Directed Mutagenesis kit (Agilent, USA) in both pMDC32 and pSITE (Huogen *et al*., 2022). *A. tumefaciens* (GV3101 strain) was subsequently transformed with each construct. The viral RNA silencing suppressor protein P19 was cloned earlier in pMDC32 (Chu *et al*., 2000; Bera *et al*., 2022)..

### Expression of constructs in host plants

For stable expression, wild type Arabidopsis plants (Col-0 strain) were transformed with the empty plasmid vector pMDC32 (EV), pMDC32 NIa-Pro, or the pMDC32 NIa-Pro protease mutant C151A using the *A. tumefaciens* above and the floral dip method (Zhang *et al*., 2006). The seeds of the MED16-FLAG complemented lines were generously provided by Dr. Zhonglin Mou at University of Florida, USA (Wang *et al*., 2015).

For transient expression of NIa-Pro, four-week-old *N. benthamiana* plants were infiltrated with overnight bacterial cultures of *A. tumefaciens* GV3101 cells carrying the pSITE empty vector, pSITE:NIa-Pro, or pSITE:NIa-Pro C151A mutant along with the pMDC32:P19 silencing suppressor. Bacteria were resuspended in 0.1M magnesium sulfate and 150µM acetyl syringone to obtain a final OD_600_ of 0.2. Plants were infiltrated with a needle-less syringe as described above.

To evaluate MED16 cleavage by NIa-Pro, the MED16-FLAG complemented plants were crossed with the Arabidopsis plants overexpressing the empty plasmid vector pMDC32, pMDC32 NIa-Pro, or the pMDC32 NIa-Pro (C151A). Pollen from the pMDC32 EV, NIa-Pro, or NIa-Pro C151 lines were used to pollinate MED16-FLAG plants. The F1 progeny were verified for the presence of NIa-Pro with RT-PCR as described below in the RT-PCR section. F1 and subsequent F2 plants were self-crossed to obtain homozygous F3 progeny carrying MED-FLAG with pMDC32-EV, pMDC32-NIa-Pro, and pMDC32-NIa-Pro C151A mutant that were used to collect tissue for total protein extraction and nuclear separation.

### Aphid fecundity bioassays

For stable and transient expression experiments, one adult *M. persicae* was placed on a single leaf of each transgenic Arabidopsis plant, or on each agrobacterium-infiltrated *N. benthamiana* leaf 24 hours after infiltration, using cages. Twenty-four hours later, all but one nymph was removed from the plant along with the adult aphid. The single nymph was allowed to develop on each plant or leaf for nine days for Arabidopsis and seven days for *N. benthamiana*, and then the number of aphids were counted. This was adequate time for the nymph to reach adulthood on each plant and to start producing their own nymphs. A similar set-up was used for bioassays with wild type and the *med16* mutant plants (see methods below). For all fecundity experiments at least 12 separate plants were used per treatment for each replicate, and each experiment was replicated at least twice.

### Screening the Arabidopsis transcriptome for NIa-Pro cleavage sites, MED16 nuclear localization signals (NLS), and MED16 isoforms

Recently we characterized the transcriptome of Arabidopsis plants with and without TuMV infection and aphid-infestation (Bera *et al*., 2022). To identify putative host proteins that are potentially cleaved by NIa-Pro, protein sequences were obtained for all differentially expressed transcript using the Arabidopsis proteome uniprot-proteome_UP000006548. Protein sequences were searched using the Perl *grep()* function with the search term corresponding to the most common NIa-Pro cleavage sequence (VxxQ), while the MED16 protein sequences were searched manually using the find function in Microsoft Word for the second most common cleavage sequence (VxxE) (Adams *et al*., 2005). We used the NLStradamus tool from University of Toronto to locate the nuclear localization signal of the MED16 protein sequences from Arabidopsis and *N. benthamiana.* We used Kallisto version 0.48.0 for the quantification of MED16 isoforms (Bray et al. 2016). First, a Kallisto index was constructed using the TAIR10 full-length cDNA obtained from Ensembl Plants (https://plants.ensembl.org/) release 107. Trimmed reads generated from each sample were pseudo-aligned to this index with 100 bootstraps using the following command: ‘kallisto quant - b 100 --single -l 200 -s 20’ to produce an abundance table with transcripts per million (TPM) values for each sample. We used I-TASSER for protein structure predictions of the different MED16 isoforms and for solvent accessibility scores of the different NIa-Pro cleavage sites (Zhou et al., 2019; Zhou et al., 2022).

### MED16 transcript abundance and cleavage experiment set-up

Three-week-old Col-0 Arabidopsis plants were rub-inoculated with TuMV or mock-inoculated as described above. One week after rub-inoculations, virus infected plants were identified by using UV light to visualize the GFP. One cage was added to a single fully infected leaf for six TuMV-infected plants, or to a single developmentally matched leaf from six mock-inoculated plants. Next, 20 adult aphids were added to each cage. As a control, cages were added without aphids to six additional plants for both treatments (virus- and mock-inoculated). The exact same methods were used to set-up aphid and no aphid treatments on four-week-old Arabidopsis plants overexpressing pMDC32 EV, pMDC32:NIa-Pro, or pMDC32:NIa-Pro C151A protease mutant(N = 3-7 plants per treatment). Six to seven plants each were used for pMDC32 EV and pMDC32:NIa-Pro C151A with or without aphids, while four plants were used for pMDC32:NIa-Pro and three plants were used for pMDC32:NIa-Pro that were infested with aphids. For western blot analyses, plant tissue was collected from four-week-old Arabidopsis plants with or without aphid feeding that were overexpressing MED16-FLAG X pMDC32:EV, MED16-FLAG X pMDC32:NIa-Pro, or MED16-FLAG X pMDC32:NIa-Pro C151A mutant (six plants per treatment); N =6).

All cages and aphids were removed 48 hours after infestation. Immediately after aphid and cage removal, tissue was collected, flash frozen in liquid nitrogen, and stored at −80°C. For RNA extractions, 100mg of leaf tissue was collected separately from each plant replicate for each treatment. For total protein extractions, 50mg of tissue was collected from each plant for a total of 0.3g of pooled tissue per treatment. Similarly, another 0.3g of tissue was pooled for each of the treatment to perform nuclear protein extraction and separation.

### RNA extraction, cDNA synthesis, RT-PCR, and quantitative RT-PCR

Tissue was homogenized using liquid nitrogen and one metal bead of 1/8 inch diameter and shaken in a 5-gallon Harbil paint shaker (Fluid Management Inc, Wheeling, USA) for 20 secs three times while immersing tissue vials in liquid nitrogen between homogenization to prevent the tissue from thawing. Total RNA was extracted using the Quick-RNA kit (Zymo Research, CA, <u>USA</u>). One microgram of total RNA was used to synthesize cDNA using Oligo dT primers and Smart MMLV Reverse Transcriptase kit (Takara Bio Inc, Shiga, Japan). The expression of NIa-Pro was verified in all samples using gene specific primers and RT-PCR (Fig S1a, Table S1). For quantitative RT-PCR (qRT-PCR), cDNA was diluted 10 fold in nuclease free water and reactions were performed in a Bio-Rad CFX96 system using gene specific primers (Table S1) and Bio-Rad SYBR green master mix (Bio-Rad, Hercules, USA). Each sample was run in triplicate. Cq values were exported and averaged across technical replicates for each sample and then relative expression of all treatments was calculated using the 2^−ΔΔCt^ method (Livak and Schmittgen, 2001). *UBIQUITIN* was used as the endogenous transcript control for the target genes *MED16, VSP2* and *PDF1.2,* and the undamaged mock-infested or undamaged empty vector lines were used as the treatment controls.

The presence of NIa-Pro in virus-infected plants, pmDC32-NIa-Pro and pMDC32-NIa-Pro C151A overexpression plants, and MED16-3FLAG plants crossed with EV, pmDC32-NIa-Pro and pMDC32-NIa-Pro C151A overexpression plants was verified using sequence specific primers sets (Table S1; Fig. S1). RNA was extracted from these plants and cDNA was made as described above which was then used as a template, and RT-PCR was performed using Go-Taq polymerase (Promega, WI, USA).

### Total protein extractions and immunoblot assays

Plant tissue was homogenized in a mortar and pestle with liquid nitrogen and total protein extracted in 3ml of ice-cold lysis buffer containing 0.5M sodium citrate, 5% sodium dodecyl sulfate, 1.5M sodium chloride, 2% ß-mercaptoethanol and 2 tablets/10ml of complete EDTA free protease inhibitor (Sigma Aldrich, USA), 100μM E-64 cysteine protease inhibitor (Bera *et al*., 2022). Samples were then boiled at 100°C for 10mins followed by centrifugation at 14000g at room temperature for 10mins in a tabletop centrifuge (Eppendorf 5425, Hamburg, Germany). The supernatant was collected and used to quantify total extracted protein using a BCA protein quantification kit compatible with denaturing agents such as SDS and ß-mercaptoethanol (Thermo Fisher Scientific, Whatham, USA). Fifty micrograms of protein for each treatment was loaded onto a (4-20)% acrylamide SDS-PAGE denaturing protein gel (Bio-Rad, Hercules, USA) and separated at 4°C at 150 constant volts. Proteins were then transferred to a nitrocellulose membrane and blocked with 5% milk in 1x Tris buffered saline (TBS) for 2hr. The blots were then incubated in antibodies overnight at 4°C in 5% milk in 1x Tris buffered saline. To visualize MED16-FLAG protein, a FLAG-HRP antibody from Mittenyl Biotech (Bergisch Gladbach, Germany) were used at 1:5000 dilution. Blots were visualized using the SuperSignal West Femto Maximum Sensitivity Substrate chemiluminescence kit (Thermo Scientific, Whatham, USA) on a Chemidoc (Biorad, Hercules, USA).

### Separation of nucleus and cytosol localized proteins and immunoblots

Tissue was homogenized in a mortar and pestle with liquid nitrogen and 3ml of ice-cold lysis buffer was added consisting of 20mM Tris-HCl (pH=7.4), 25% glycerol, 2mM EDTA, 25mM MgCl_2_, 40mM KCl, 250mM sucrose, 100μM E-64 and 1mM DTT (Zhao *et al*., 2022). The lysate was filtered through 100-micron and 40-micron filters sequentially on ice. The filtered lysate was centrifuged at 1500g for 10min at 4°C, and the supernatant was stored as the cytosolic fraction. The pellet consisting of the nuclear fraction was resuspended in nuclear resuspension buffer consisting of 20mM Tris-HCl (pH=7.4), 25% glycerol, 25mM MgCl_2_, 100μM E-64 and 0.1% Tween-20 and centrifuged for 1500g for 10min at 4°C to wash the nuclear fraction. The nuclear fraction was washed five times and then the nuclear pellet was weighed, and the pellet was resuspended in the nuclear resuspension buffer described above. The nuclear and cytosolic fractions were boiled in 1X Laemmli buffer (Laemmli, 1970) for 10mins at 100°C and protein was quantified using BCA compatible protein kit (Thermo Fisher Scientific, Whatham, USA). Fifty micrograms of protein were loaded onto a 4-20% gradient SDS-PAGE denaturing protein gel as described above, and immunoblot detection of MED16 protein was performed as described above using the anti-FLAG antibody. To evaluate successful nuclear and cytosolic protein separations, histone H3 and phosphoenol pyruvate carboxylase protein primary antibodies were used respectively in the ratio 1:10000 (Agrisera biotech, Sweden). Histone proteins are localized only to nucleus, while phosphenol pyruvate carboxylase is an enzyme that is involved in carbon metabolism in plants and is present only in cytosol (Sonawane *et al*., 2018; Talbert *et al*., 2002). Anti-rabbit secondary antibody conjugated with horse radish peroxidase enzyme was used in the ratio of 1:10000 (Sigma Aldrich, USA) and blots were visualized using chemiluminescent substrate as described above.

### Aphid fecundity and TuMV infection bioassays on *med16* mutants

One adult *M. persicae* was placed on a single leaf of wildtype Col-0 and *med16* mutant Arabidopsis plants using cages. Twenty-four hours later, all aphids were removed except for one nymph from each cage. The single nymph per cage was allowed to develop on for nine days and then the number of progeny and the founder counted. For TuMV infection bioassays, a single leaf of wildtype Col-0 and *med16* mutant Arabidopsis was inoculated with TuMV as described above. Five days later, the number of plants with GFP visible in the rosette was counted. Ten separate plants of Col-0 and twelve plants of *med16* mutant, were used for evaluating aphid fecundity while 30 plants of each genotype were used to evaluate TuMV infection, and both experiments were replicated twice.

### Data analysis

Fecundity was analyzed by one-way ANOVA using the aov function in R (version RStudio 2022.07.2) at *p* value ≤ 0.05. Means separation was done using Tukey HSD post-hoc test using the TukeyHSD function in R. Two-way ANOVA tests were performed for gene expression data with the relative quantification (RQ) with insect presence and the plant treatment as the dependent variables. Multiple comparisons of means were done using post-hoc Tukey test using the emmeans function in R.

## Results

### The protease activity of viral protein NIa-Pro is required for increasing aphid performance on plants

Transient expression of the viral protease NIa-Pro has been shown to inhibit plant defenses that target aphids and increase aphid performance (Casteel *et al*., 2014). To determine if the protease activity of NIa-Pro is required for this phenotype, we mutated the active site cysteine (C151) to alanine (C151A) to create a protease inactive version of the protein as previously described (Huogen *et al*., 2022). Consistent with our previous studies (Casteel *et al*., 2014), expression of NIa-Pro in Arabidopsis (Fig 1a; F_2,35_= 7.74, p = 0.00165) and *N. benthamiana* (Fig 1b; F_2,32_= 43.82, p<0.0001) increased aphid performance compared to plants expressing the empty expression vector. In contrast, expression of the protease dead NIa-Pro (C151A) did not significantly enhance aphid fecundity compared to the controls in either host plants (Fig 1a, b). Taken together this suggests that the protease activity of NIa-Pro is required for its ability to inhibit plant defenses and increase aphid performance.

**Fig 1.**
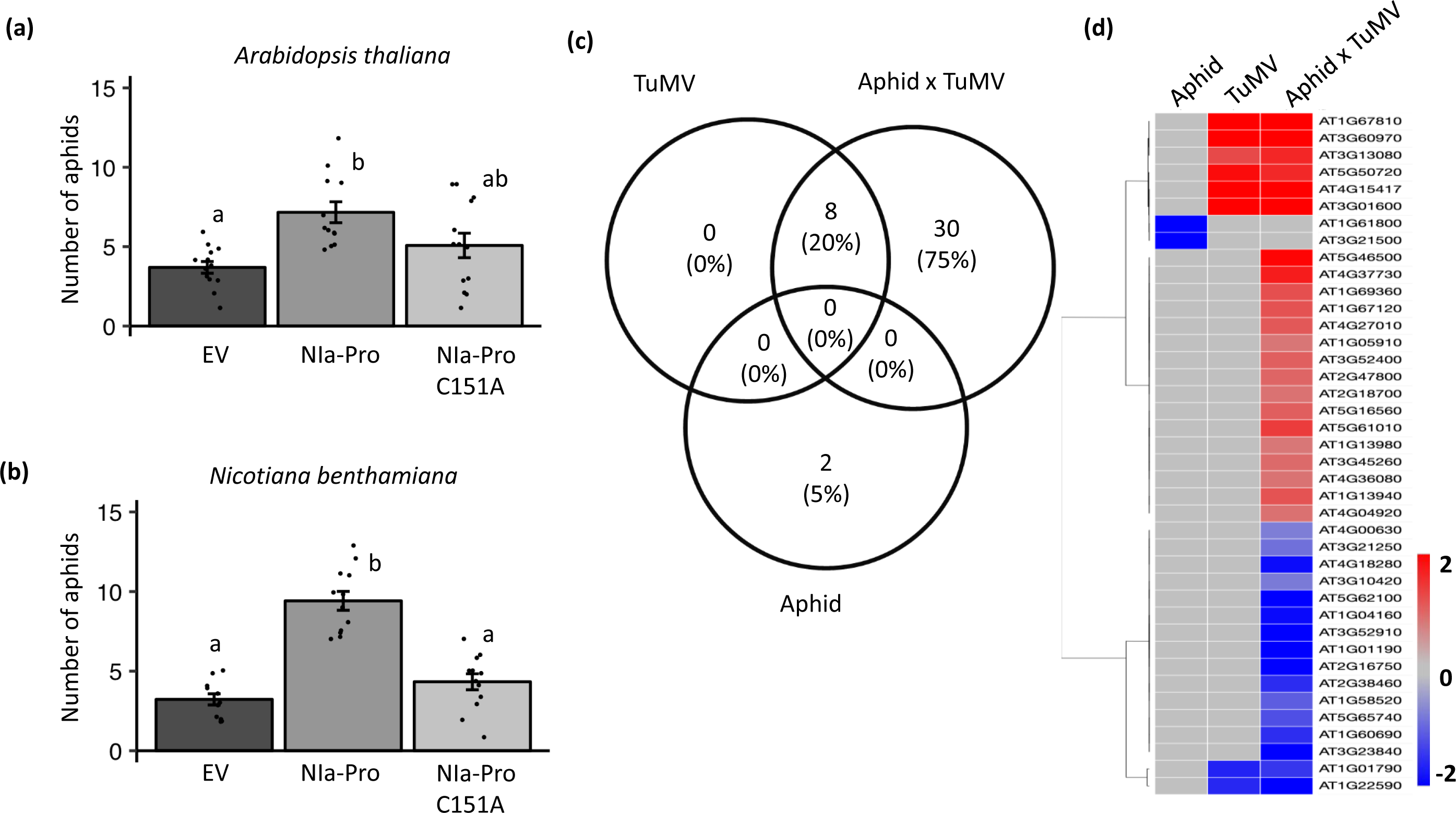
Protease activity of the viral protein NIa-Pro is required to increase aphid performance in plants. The number of progeny produced by a single one-day old *Myzus persica*e foundress after (a) 9 days on *Arabidopsis thaliana* or (b) 8 days on *Nicotiana benthamiana* plants overexpressing an empty expression plasmid (EV), NIa-Pro, or the NIa-Pro mutant (C151A) that lacks protease activity. Aphid reproduction was higher on plants expressing NIa-Pro compared to the EV control, while there was no difference in aphid progeny when feeding on NIa-Pro C151A compared to EV controls. (c)The number transcripts encoding proteins with predicted NIa-Pro cleavage sites (V-X-H-Q.) from plants infected with turnip mosaic virus (TuMV) and/or fed upon by *M. persicae* (aphid) compared to mock-inoculated, aphid-free plants. (d) A heatmap showing the up- and down-regulation of the same set of transcripts from (c). Data in (a) and (b) were analyzed using one-way ANOVAs and means separation was performed using Tukey HSD test. Letters represent significant differences among treatments using a p*-*value < 0.05; error bars indicate standard error; *N* = 12 for (a) and (b).

### Aphid- and virus-induced Arabidopsis transcripts encode proteins with NIa-Pro cleavage sites

Proteases from animal viruses can cleave host proteins with specific cleavage sequences to regulate host defenses (Chin and Saeed, 2022; Lawrence *et al*., 2012). To investigate if the potyviral protease NIa-Pro could cleave plant proteins that are required for plant defenses, we searched for the most common NIa-Pro cleavage site (VxHQ) in the protein sequences of the TuMV and/or *M. persicae* regulated Arabidopsis genes from a recent transcriptome study (Table 1; Bera et al. 2023). Only 40 of the 1626 DEGs contained NIa-Pro cleavage sites in their predicted coding sequence across all treatments (Table 1). The mRNA transcripts of most of the predicted plant proteins with NIa-Pro cleavage sites were differentially expressed in plants infected with TuMV and fed on by aphids (38 DEGs, Table 1, TuMV x Aphid column; Fig. 1c). Roughly 50% of the 38 DEGs were up-regulated and down-regulated in response to TuMV infection and aphid feeding compared to controls (Fig. 1d; 16 down, 22 up, Table 1, TuMV x Aphid column). Eight of the DEGs in plants infected with TuMV alone contained predicted NIa-Pro cleavage (Table 1; Fig. 1c). All eight of these were also differentially regulated in plants infected with TuMV and fed upon by aphids (Table 1, TuMV column & TuMV x Aphids; Fig. 1d), and they were regulated in the same direction and magnitude in both treatments, suggesting these are not impacted by aphid presence (Lawrence *et al*., 2012 Fig. 1d; Table 1, TuMV & TuMV x Aphid columns). Only two of the DEGs in response to aphid feeding alone encoded proteins with predicted NIa-Pro cleavage sites, and both were down-regulated compared to controls (Fig. 1c, d; Table 1, Aphid column).

**Table 1:**
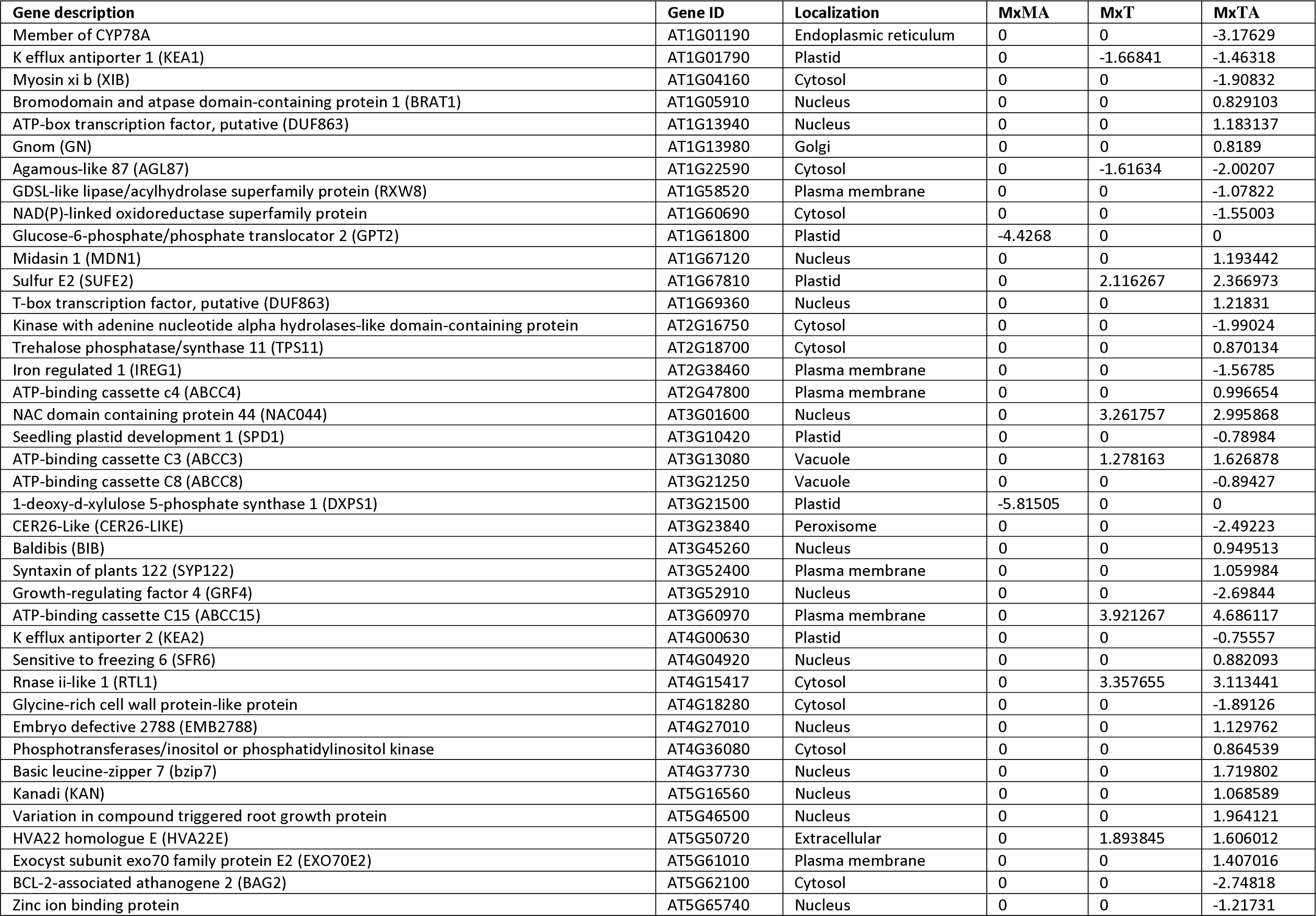
List of genes that are induced by virus infection in Arabidopsis and has potential cleavage site for NIa-Pro.

We searched the 40 candidates for genes already known to be involved in plant defense responses against aphids or virus infection. Among the genes, we found SYP122, a Qa-SNARE protein involved in callose deposition, vesicle fusion (Rubiato *et al*., 2022), and a known negative regulator of programmed cell death, salicylic acid signaling, and jasmonic acid signaling (Zhang *et al*., 2007). Recently, it was also shown that SYP122 transcripts are induced by *M. persicae* feeding on Arabidopsis plants (Annacondia *et al*., 2021). TREHALOSE PHOSPHATASE SYNTHASE 11 (TPS11), a protein shown previously to induce resistance to aphids in Arabidopsis (Singh *et al*., 2011), and RNASE-II like 1 (RTL1), a protein that has been shown previously to suppress RNA silencing in host plants (Sehki *et al*., 2023), were two other defense related genes found among the candidates.

A particularly interesting defense related candidate for us was SENSITIVE FOR FREEZING 6 (SFR6) or MEDIATOR 16 (ATG404920), a plant protein involved in regulating jasmonic acid and ethylene dependent defense responses and resistance to necrotrophic pathogens (Wang *et al*., 2015). MED16 has also been shown to bind to the transcription factor WRKY33 (Zhang *et al*., 2012; Wang *et al*., 2015), an important negative regulator of ABA and SA responses, and activator of ET dependent plant defenses (Liu *et al*., 2015). Previously, we demonstrated that aphid herbivory induces ethylene-dependent plant defenses that are inhibited in potyvirus-infected plants (Casteel *et al*., 2015; Bak *et al*., 2019). This led us to hypothesize that potyvirus inhibition of ethylene dependent defenses could be due to MED16 cleavage by NIa-Pro, and this could explain why aphid fecundity was lower in plants that expressed NIa-Pro that did not have protease activity (Fig. 1a, b).

### The MED16 protein has multiple NIa-Pro cleavage sites upstream of a nuclear localization signal

To determine if NIa-Pro cleaves MED16 we conducted immunoblots with *med16* mutant Arabidopsis that have been complemented with a FLAG tagged version of MED16 with and without TuMV infection. In immunoblots, we detected two bands between 180 and 130 kD in all treatments, which corresponds to the size of MED16 and is consistent with previous studies (Fig. 2a) (Wang *et al*., 2015). We also detected a ∼36kD protein product in the virus treatment, which was barely detected in the controls (Fig. 2a; black arrow).

**Fig 2.**
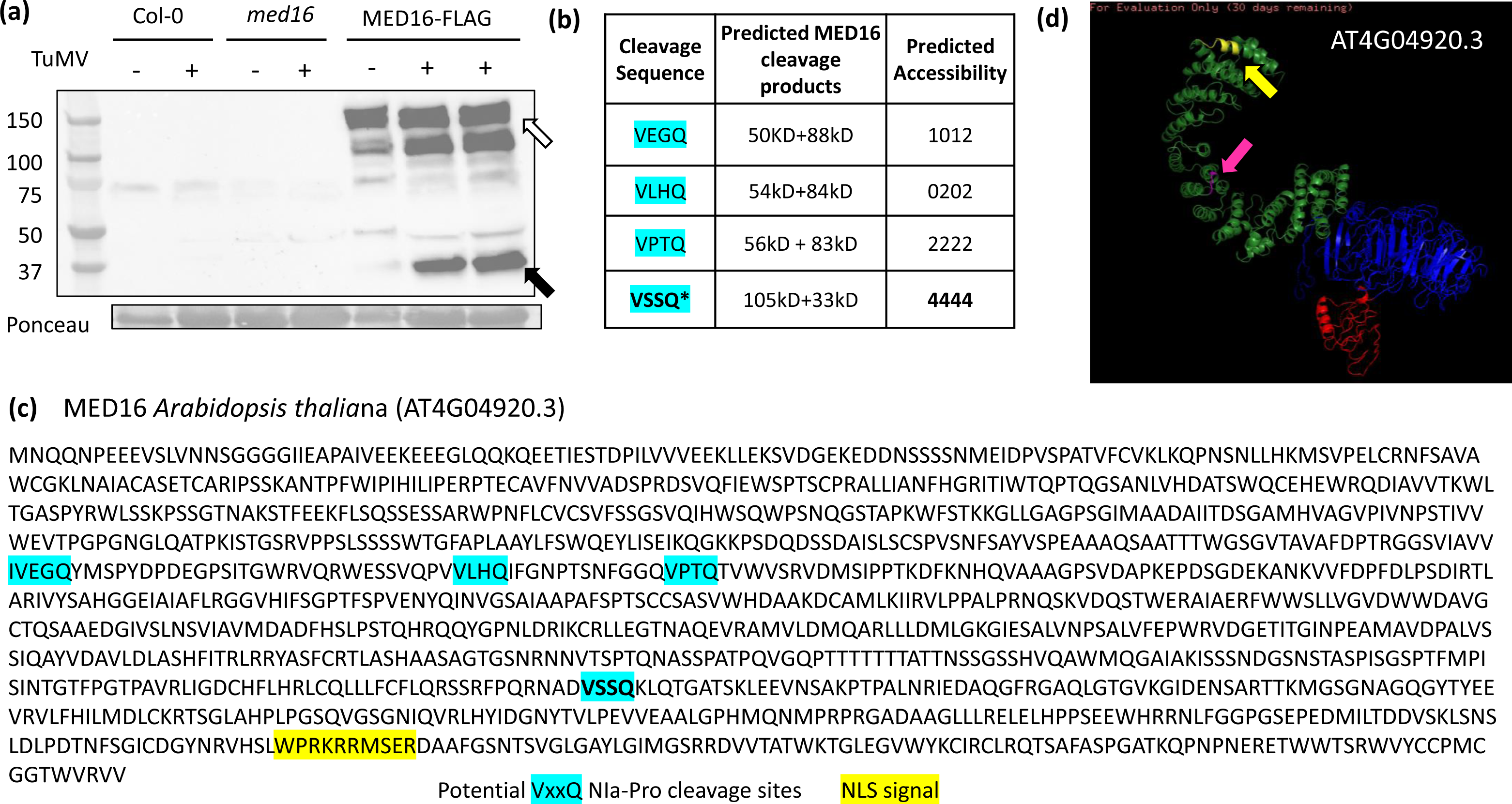
The MEDIATOR16 (MED16) protein is cleaved upstream of a nuclear localization signal (NLS) in the presence of turnip mosaic virus. (a) Col-0 wildtype Arabidopsis, *med16* mutants, and *med16* mutants complemented with MED16-3XFLAG were mock-inoculated or inoculated with turnip mosaic virus and then protein abundance of *MED16* was measured using immunoblots and FLAG antibodies. The full length MED16 protein (white arrows) was detected as well as a smaller ∼36kDa cleaved product (black arrows). Each lane represents proteins extracted from 5 individual plants. Equal amounts of protein were loaded into each well as shown by Ponceau stain. The Arabidopsis MED16 protein sequence was searched for the most common NIa-Pro cleavage site (VxxQ). Predicted cleavage product sizes and amino acid predicted solvent accessibility scores are shown in (b). (c) The location of the NIa-Pro cleavage sites and of the nuclear localization signal (NLS) in the amino acid sequence are presented. (d) The NLS (yellow arrow) and the VSSQ cleavage site (pink arrow) locations are presented in the predicted protein structure of AT4G04920.3).

To investigate if this could be a cleavage product of NIa-Pro, we searched for the location of the Arabidopsis MED16 VxxQ cleavage sequences (Goh and Hahn, 2021). We found three predicted isoforms of MED16 (Fig. S3), and each isoform sequence contained four potential VxxQ cleavage sites (Fig. 2b,c). We determined AT4G04920.3 was the most abundant isoform of MED16 (Fig. S4a) by reanalyzing an Arabidopsis transcriptome data set we recently published (Bera *et al*. 2021). To determine if all four predicted NIa-Pro cleavage sites were accessible we used I-TASSER to predict the structure of the most abundant MED16 isoform (Fig. 2d), as well as the other two isoforms (Fig S4b). One site, VSSQ, was predicted to be more accessible than the other sites (Fig. 2b), and to produce a cleavage product around ∼36kDA with the 3kDA FLAG tag, which corresponded to the immunoblot (Fig. 2a). When comparing the predicted structures of the different isoforms, we noticed an open hook like structure for AT4G04920.3, the most abundant form, that was not present for the other two isoforms (Fig. 2d; Fig. S4b). Notably, aphid feeding reduced the abundance of this form of MED16 (Fig. S4a), however the functional relevance of this was not determined. To determine how cleavage could impact function, we next examined MED16 for functional motifs and localization signals and found a nuclear localization signal (NLS) present downstream of the potential NIa-Pro cleavage sites (Fig. 2c). These results suggest NIa-Pro may cleave off the NLS from MED16 and prevent it from localizing to the nucleus, where it normally functions.

### Presence of aphids and virus increased MED16 transcripts, protein levels, and cleavage

We next evaluated the impact of aphid feeding and virus infection on *MED16* mRNA abundance in an independent experiment. Virus infection significantly increased *MED16* transcripts compared to controls (Fig. 3a; Main effect, virus: F_1,22_ = 18.007, p-value = 0.0003), however there were no significant impacts of aphid feeding or interactions between aphid and virus on *MED16* transcript accumulation (Fig. 3a; Main effect, aphid: F_1,22_ = 0.680, p-value = 0.418; virus x aphid: F_1,22_= 0.262, p-value = 0.614). Upon performing means separation with a Tukey test between the treatments, we observe that *MED16* transcripts were higher in Arabidopsis plants at 10 days post TuMV infection (2.71 +/- 0.444; Fig. 3a) and in virus infected-plants infested with aphids (3.41 +/- 0.819; Fig. 3a), compared to mock-inoculated plants (0.686 +/- 0.391; Fig. 3a) or mock-inoculated plants infested with aphids (0.833 +/- 0.167; Fig. 3a). Notably, immunodetection showed MED16 protein abundance increased in virus-infected plants and in plants with aphids feeding compared to the mock controls (Fig. 3b). We also detected the ∼36kDa product in the aphid and virus treatment, however it was barely detected in virus infected plant alone (Fig. 3b). Together these results suggest MED16 is induced in the presence of aphids or TuMV, and protein cleavage is increased when both challengers are present.

**Fig 3:**
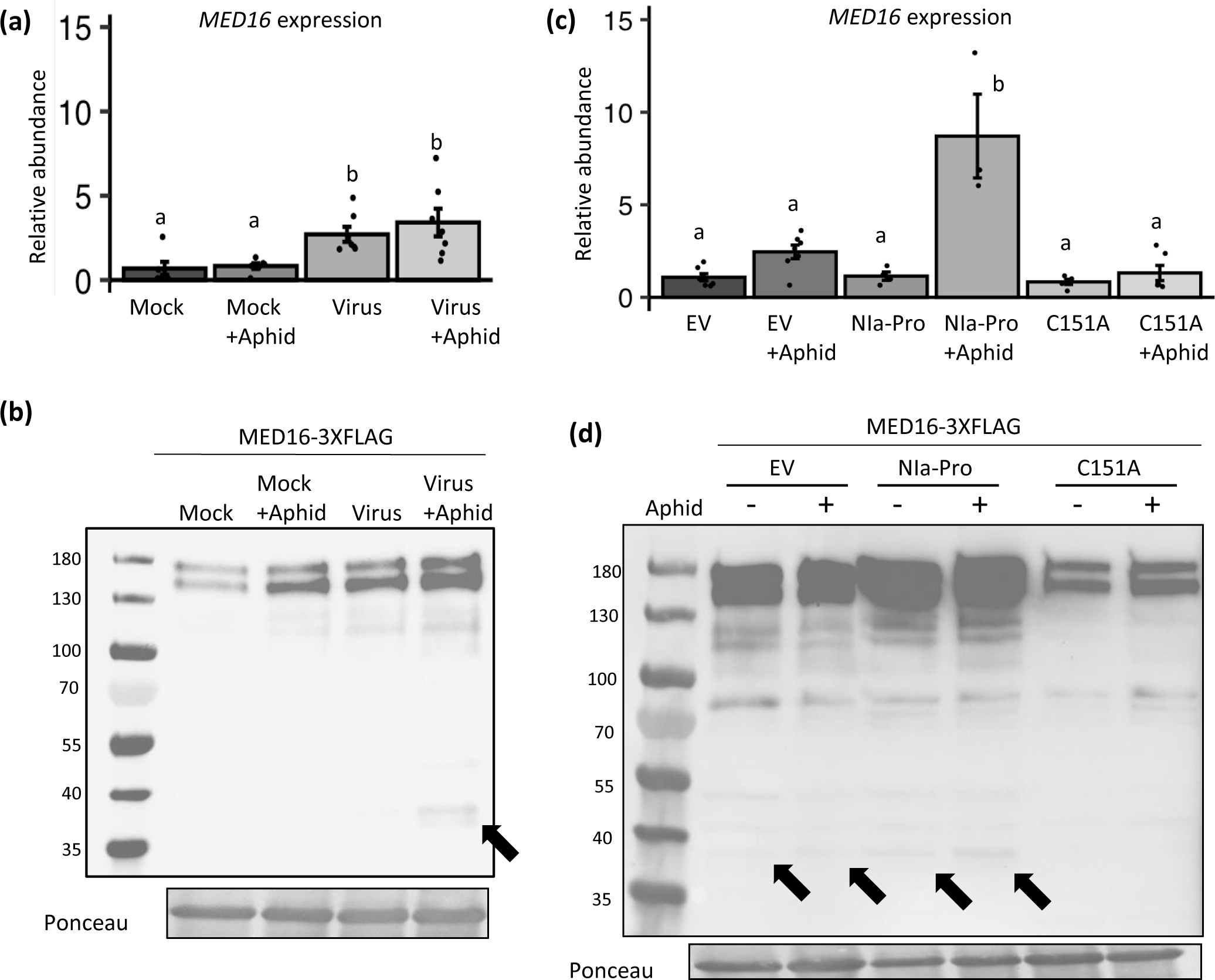
*MEDIATOR16* transcripts and proteins are induced in the presence of turnip mosaic virus and the viral protein NIa-Pro. Relative expression of *MED16* in (a) mock- and virus-infected plants with and without aphid feeding, and in (b) plants overexpressing the empty expression plasmid (EV), NIa-Pro, or the NIa-Pro (C151A) mutant that lacks protease activity, with and without aphid feeding. (c, d) Protein abundance of *MED16* was measured in *med16* mutants that were complemented with MED16-3XFLAG using immunoblots and *FLAG* antibodies. In (c) complemented plants were mock- and virus-infected with and without aphid feeding, and in (d) complemented plants were crossed with plants overexpressing NIa-Pro, EV, or NIa-Pro C151A mutant. The full length MED16 protein (140kD) as well as smaller ∼36kDa cleaved product (black arrows) were detected. In (c, d) each lane represents proteins extracted from 5 individual plants. Equal amounts of protein were loaded into each well as shown by Ponceau stain. For (a, b) *UBIQUITIN* was used as an endogenous control, while mock or EV treatments were used as the treatment controls for calculating relative expression. qRT-PCR data were analyzed using two-way ANOVAs and means separation was performed using Tukey test (N = 5 −7 for a, and N = 3 - 7 for b). Bars with different letters indicate significant differences at p-value < 0.05; error bars indicate standard error.

### NIa-Pro and aphid presence enhance MED16 levels and cleavage

We next evaluated if NIa-Pro alone can increase MED16 transcript and protein levels using transgenic NIa-Pro plants that had been crossed with the MED16-FLAG complemented *med16* mutants. In contrast to the TuMV experiments above, aphid infestation increased *MED16* mRNAs (Main effect, aphid: F_2,26_ = 16.92, p-value < 0.0001; Fig 3c). After performing a means separation using Tukey test at we observed that Arabidopsis plants expressing NIa-Pro in presence of aphid feeding had significantly higher levels of *MED16* transcripts (8.71 +/- 2.26) than all other treatments, including the plants expressing the NIa-Pro protease mutant (C151A) with aphid feeding (1.32 +/- 0.409) or without aphid feeding (0.834 +/- 0.139) (plant genotype x aphid: F_2,26_ = 17.64, p-value < 0.0001; Fig. 3c). While MED16 transcript levels were not elevated in plants expressing NIa-Pro without aphids compared to the controls (Fig. 3c), MED16 protein levels were elevated in Arabidopsis expressing NIa-Pro with or without aphids, and much greater than all other treatments (Fig. 3d). In this experiment, the ∼36kD protein product was detected in all treatments (Fig. 3d; black arrows), except for the plants expressing the NIa-Pro protease mutant (C151A) with or without aphid feeding (Fig. 3d; black arrows). Overall, the highest amount of MED16 and of the ∼36kD cleaved protein product was observed in the plants expressing NIa-Pro in presence of aphid feeding, while MED16 levels were the lowest for plants expressing the NIa-Pro C151A mutant with or without aphid feeding (Fig. 3d). This shows that NIa-Pro and aphid presence increases MED16 levels and cleavage, and that NIa-Pro’s protease activity is required for such increases.

### Aphid feeding in the presence of TuMV or NIa-Pro increased MED16 nuclear localization and cytosolic cleavage

As a transcription regulator, MED16’s primary functions occur in the nucleus. Virus infection increased the accumulation of the full length MED16 in the nucleus (Fig. 4a), and aphid feeding enhanced MED16 nuclear accumulation even more (Fig. 4a). Likewise, accumulation of the full length MED16 increased in plants expressing NIa-Pro, and this was enhanced in plants that had been fed on by aphids (Fig. 4b). To our surprise aphid feeding alone reduced MED16 accumulation in the nucleus for empty vector controls (Fig. 4b). Notably, the ∼36 kD product was not detected in any nuclear fractions (Fig. 4b), however protein levels in general were much lower.

**Fig 4:**
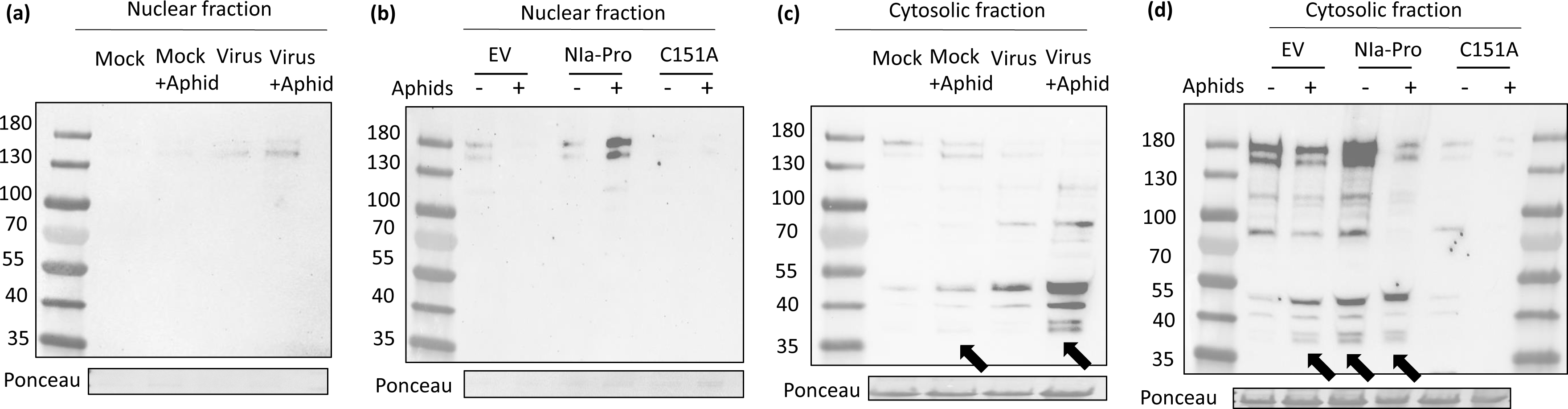
Aphid feeding in the presence of TuMV or NIa-Pro increased MED16 nuclear localization, and cytosolic cleavage. Immunoblot detection of the MED16 protein in the (a, b) nuclear and (c, d) cytosolic fractions of MED16-3XFLAG complemented Arabidopsis plants. (a, c) Plants infected with turnip mosaic virus with and without aphid feeding and (b, d) plants overexpressing the empty vector (EV), NIa-Pro, or the protease mutant NIa-Pro (C151A), with and without aphid feeding. The ∼36kDA cleaved product of MED16 is indicated by black arrows. Each lane represents proteins extracted from 6 individual plants. An equal amount of protein was loaded in each well as shown by Ponceau stain, and MED16 was detected using a FLAG antibody.

Our previous results demonstrated that NIa-Pro is located inside the nucleus and cytosol, however it must be outside of the nucleus to inhibit plant defenses and enhance aphid fecundity (Bak *et al*., 2017). This led us to hypothesize that NIa-Pro may cleave MED16 in the cytosol before it is able to relocate to the nucleus. To address this hypothesis, we next measured the amount of full length MED16 and MED16 cleavage products in the cytosol. Virus infection alone and in combination with aphid feeding reduced accumulation of full length MED16 in the cytosol (Fig. 4c). While the full length MED16 was barely present in the cytosol of plants with aphid and virus feeding, there was a significant increase in MED16 cleaved products in the cytosol, including accumulation of the ∼36 kD product (Fig. 4c). Notably, low levels of the ∼36 kD product were only detected in the cytosol of the mock-inoculated plants with aphid feeding (Fig. 4c).

As with the virus infection, accumulation of the full length MED16 in the cytosol was the lowest for the NIa-Pro plants with aphid feeding (Fig. 4d). Aphid feeding alone also reduced the accumulation of full length MED16 in the cytosol, while NIa-Pro expression enhanced it (Fig. 4d), a pattern that was also observed in the nucleus (Fig. 4b). Despite the fact that only NIa-Pro expression in the presence of aphids had a large impact on the accumulation of full length MED16, accumulation of the ∼36 kD product increased in all three aphid and NIa-Pro treatments (Fig. 4d). Full length MED16 and cleavage products were also measured in the nucleus and cytosol of transgenic plants expressing the NIa-Pro C151A mutant with and without aphid feeding (Fig. 4b, d). Levels of full length MED16 were extremely low in the nucleus in the NIa-Pro C151A plants regardless of aphid presence (Fig. 4b), and similar to levels in aphid control plants (EV + aphids; Fig. 4b). Notably, the ∼36 kD MED16 product was not detected in either the nucleus or the cytosol for any of the NIa-Pro protease mutant treatments (Fig. 4c, d). In combination, these results suggest that aphids and NIa-Pro induce MED16 cleavage in the cytosol, potentially reducing the amount of MED16 that can relocate to the nucleus and activate defense.

### TuMV infection and NIa-Pro expression suppress MEDIATOR16-dependent transcript accumulation

If NIa-Pro MED16 cleavage reduces the pool of active MED16 in the nucleus, we hypothesized that MED16-dependent gene expression would also be reduced. To address this hypothesis, we measured transcript abundance of *PLANT DEFENSIN 1.2* (*PDF1.2*), which has been shown previously to be MED16-dependent (Wang *et al*., 2015), and the abundance of a MED16-independent transcript, JA-dependent *VEGETATATIVE STORAGE PROTEIN2* or *VSP2* (Lorenzo *et al*., 2003). The expression of *PDF1.2 transcript* was suppressed overall in presence of the virus when we use presence or absence of virus as the main effect in our two-way ANOVA model regardless of aphid feeding (main effect, virus: F_1,22_ = 6.493, p-value = 0.0183). On the other hand, aphid herbivory had no main effect on *PDF1.2* expression (main effect, aphid: F_1,22_ = 1.590, p-value = 0.2205). There was however a marginally significant interacting effect of virus infection and aphid infestation (virus x aphid: F_1,22_ = 3.403, p-value = 0.0786). *PDF1.2* relative expression was highest in mock-inoculated plants with aphids feeding (3.92 +/- 1.59), however aphids were not able to induce *PDF1.2* expression in the virus-infected plants (0.594 +/- 0.158; Fig. 5a). This suggests that pathogen-induced defenses in plants downstream of MED16 are suppressed in virus-infected plants. Notably, aphid feeding had no main effect on the expression of *PDF1.2* in any these over-expressor lines (Fig. 5b), however there was a significant effect of the plant genotype (Fig. 5b; Main effect, plant genotype: F_1,20_ = 16.811, p- value < 0.00001) and an interactive effect of aphids and NIa-Pro on *PDF1.2* expression (Fig. 5b; aphid x plant genotype: F_1,26_ = 4.591, p-value = 0.0446). Posthoc analyses with means separation further shows that aphid feeding on EV over-expressing plant had the highest *PDF1.2* transcript levels (1.39 +/ 0.205), while Arabidopsis plants that overexpress NIa-Pro alone or with aphids had lower transcript levels for *PDF1.2* (0.341+/-0.097 and 0.340+/-0.170 respectively) compared to both mock plants alone (0.937+/-0.079) and to plants overexpressing NIa-Pro C151A mutant alone or with aphids (0.861+/-0.042 and 1.10+/-0.165 respectively) (Fig 5b). Further, *PDF1.2* transcripts levels were not significantly different from the EV controls, for the plants expressing NIa-Pro C151A (Fig. 5b).

**Fig 5.**
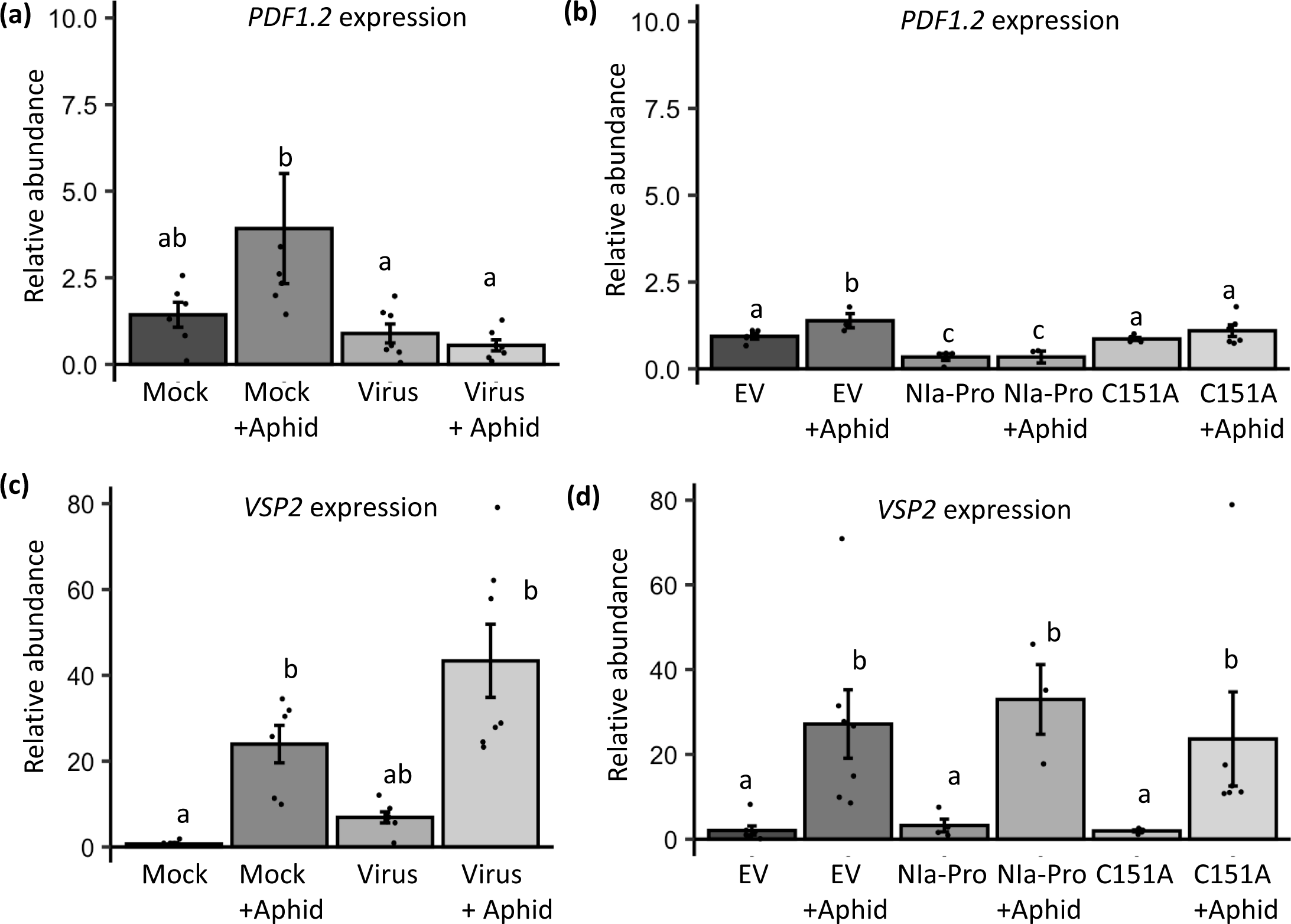
*PDF1.2* is suppressed by virus infection and the viral protein NIa-Pro, while *VSP2* expression is only induced by aphids. Relative expression of *PDF1.2* and *VSP2* was measured in (a, b) mock and virus-infected plants with and without aphids and in (c,d) plants overexpressing empty plasmid vector (EV), NIa-Pro, or the protease mutant of NIa-Pro (C151A), with and without aphids present. *UBIQUITIN* was used as an endogenous control. Mock or EV treatments were used as the treatment controls for calculating relative expression. Two-way ANOVAs were conducted and means separation performed using Tukey HSD test for all experiments (N = 5 - 7 for a & c, and N = 3 - 7 for b & d). Bars with different letters indicate significant differences at p-value < 0.05; error bars indicate standard error.

In contrast, the aphid inducible plant defense gene *VSP2,* that is not dependent on MED16, was not induced by virus infection (Fig 5c; Main effect, virus: F_1,22_ = 3.106, p-value = 0.06) and there was no interacting effect of viral infection and aphid feeding (Fig 5c; virus x aphid: F_1,22_ = 1.677, p-value = 0.209). However, *VSP2* expression was significantly induced in response to aphid feeding (Fig5c; Main effect, aphid: F_1,22_ = 35.653, p-value < 0.0001). *VSP2* expression in plants infested with aphids alone (24.0+/-4.38) and aphid-infested with virus infection (43.4+/-8.53), was significantly higher than uninfested plants (0.718 +/- 0.281) or plants with virus infestation only (6.90 +/- 1.29) (Fig 5c). Similarly, in the over-expressor lines, the only significant effect main on *VSP2* expression was from aphid infestation (Fig 5d; Main effect, aphid: F_1,26_ = 18.59, p-value = 0.0002; Main effect, plant genotype: F_1,26_ = 0.038, p-value = 0.962) or the interaction of aphid feeding and viral protein expression in plants (Fig 5d; plant genotype x aphid: F_1,26_ =0.129, p-value = 0.879). The *VSP2* gene expression in aphid fed plants of either empty vector (EV) or the NIa-Pro over-expressor lines (Fig 5d; EV = 27.2+/-8.06; NIa = 33.0+/-8.23; NIa C151A mutant = 23.6+/-11.1) were all higher than those that were not infested with aphids (Fig 5d; EV = 2.06+/-1.06; NIa = 3.21+/-1.49; NIa C151A mutant=1.95+/-0.251). This suggests that cleavage of MED16 protein by NIa-Pro does not affect components of this MED16-independent plant defense pathway.

### Performance of both virus and aphid are higher in absence of *MED16*

Next, we evaluated the percentage of plants infected with TuMV after 10 days, and the performance of aphids after 9 days of infestation, on Col-0 wild type and *med16* mutant Arabidopsis. The results showed that the percentage of *med16* plants infected with TuMV was significantly higher compared to the wild type Col-0 group (Fig. 6a; χ^2^ value = 10.17; p-value = 0.001428). Additionally, the performance of aphids was significantly higher on *med16* mutant plants compared to the wild type Col-0 group (Figure 6b; F_1,20_ = 13.87; p-value = 0.00134). These findings suggest that MED16 plays a significant role in the defense mechanisms of Arabidopsis against both TuMV and aphids.

**Fig 6.**
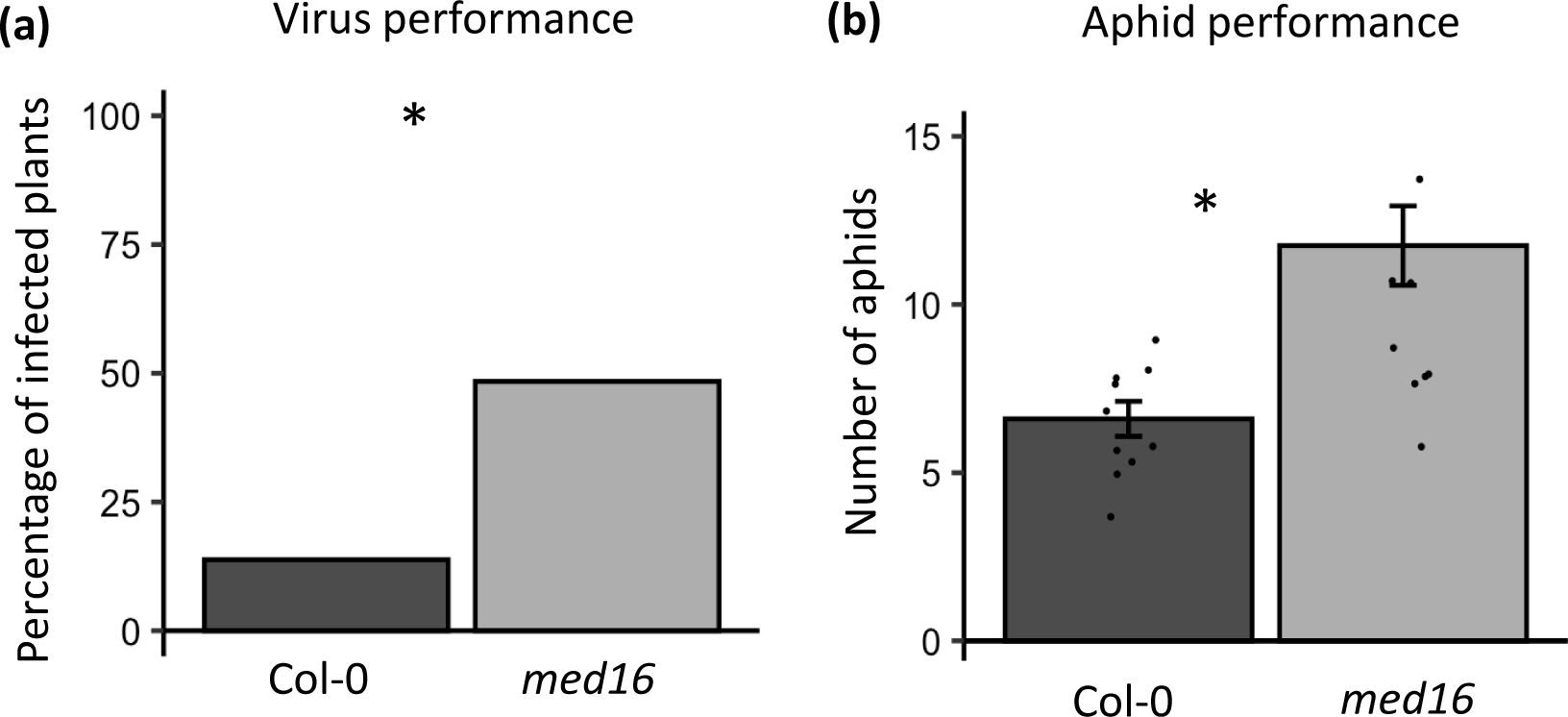
Virus and aphid performance is enhanced on *med16* mutant Arabidopsis. (a) The percentage of wild type Col-0 and *med16* mutant Arabidopsis plants infected with turnip mosaic virus (TuMV) 10 days post inoculation. (b) The number of progeny produced by a single one-day old *Myzus persica*e foundress was counted on wild type Col-0 plants and *med16* mutants after 9 days. Significant differences were determined for (a) using a *CHI-*square test. (b) Aphid data was analyzed using a one-way ANOVA and means separation was performed using Tukey HSD test. Asterisk indicates significant differences among treatments using a p- value < 0.05, and N = 30 for (a) and N = 10-12 for (b); error bars indicate standard error.

## Discussion

The viral effector NIa-Pro was previously shown to enhance aphid performance by suppressing aphid-induced plant defenses downstream of ethylene signaling pathway (Casteel *et al*., 2015; Casteel *et al*., 2014). In this study, we demonstrate that a key regulator of ethylene-dependent defenses, MED16, is cleaved in the presence of TuMV and NIa-Pro (Fig 2a & 3b), and that NIa-Pro’s protease activity is required for changes in MED16 (Fig. 3c, d), plant defenses (Fig. 5), and enhanced aphid performance (Fig. 1 a,b). NIa-Pro is a cysteine protease which contains a cysteine residue at its active site and has been shown to self-cleave itself from the viral polyprotein and then subsequently cleave other viral proteins from the polyprotein (Riechmann *et al*., 1992; Adams *et al*., 2005). In virus-infected plants, NIa-Pro is fused with another viral protein VPg which contains a NLS and accumulates in the nucleus (Schaad *et al*., 1996). In the nucleus, NIa-Pro has also been shown to also have DNase activity (Anindya and Savithri, 2004) and it has been predicted that it interacts with plant defense components previously (Rajamäki and Valkonen, 2009; Beauchemin *et al*., 2007; Riechmann *et al*., 1992).

Viral proteases have been shown to cleave host proteins in several animal systems (H., Li *et al*., 2019; Zaragoza *et al*., 2006), and more recently NIa-Pro was shown to be able to cleave plant host proteins (Huogen *et al*., 2022). In this study, over 90 proteins in the proteome of different plants (Arabidopsis, *Prunus persica* and *N. benthamiana*) were identified to have NIa-Pro cleavage sites and cleavage was verified for a few candidates (Huogen *et al*., 2022), however the biological relevance of these proteins remains largely unknown. Our work goes further by demonstrating viral and aphid performance was enhanced on *med16* mutants (Fig. 6b), and that the NIa-Pro protease activity was required for defense suppression (Fig. 5a, b). These findings provide insights into the mechanisms underlying virus-aphid-plant interactions and have implications for plant defense strategies against viral infections and insect herbivory.

In a previous study, we demonstrated that NIa-Pro re-localizes from the nucleus to the vacuole in presence of aphid feeding. When an NLS was added to NIa-Pro, it was confined completely to the nucleus and was unable to re-localize, suppress plant defenses, or increase aphid performance on host plants (Bak *et al*., 2017). This suggested that NIa-Pro’s role in plant-aphid interactions occurs outside of the nucleus, which is consistent with the current study’s findings. The nucleus contains pores that allow the entry and exit of a 50 kDa protein such as NIa-Pro (Bak *et al*., 2017). MED16, which is needed for transcription regulation and is ∼140kD is too large and requires an NLS to enter the nucleus (Knight *et al*., 2009). Indeed, a potential NLS was found at N-terminus of MED16, immediately after the NIa-Pro cleavage site (Fig. 2b, c, d). This suggests NIa-Pro inhibits MED16’s function by removing the NLS in the cytosol and preventing nuclear localization. This is further supported by the detection of the predicted ∼36 KD cleavage product only in the cytosol fractions from TuMV-infected plants and NIa-Pro expressing plants (Fig. 4 c, d). Furthermore, MED16 was only present in the nucleus as a full length protein (Fig 4 a, b).

To our knowledge, this study is the first example of a plant virus protease, and of any plant pathogen effector, inducing the removal of an NLS from a host protein to increase virulence. In animal viruses, however, it has been shown that viral proteases can remove NLS from animal proteins. For example, the viral protease 3C^pro^ of Foot-and-mouth disease virus (FMDV), can cleave the C-terminus end of host RNA-binding protein Sam68 that houses the NLS signal. The cleavage of Sam68 prevents it from being localized in the nucleus and its subsequent localization in the cytoplasm, thereby increasing the viral titer 1000 fold in fetal porcine cell lines (Lawrence *et al*., 2012). The sugar beet cyst nematode *Heterodera schachttii* secretes the 4E02 effector in host plants, which then interacts with the vacuolar plant protease RD21A (RESPONSIVE TO DEHYDRATION 21A). RD21A is a cysteine protease which moves to the nucleus when bound with 4E02 and is predicted to cleave several plant defense related proteins rendering the plants susceptible to *H. schachttii* as well as to necrotrophic pathogen *Botrytis cinerea* (Pogorelko *et al*., 2019). This work shows how a microbial effector can highjack a host protease, to relocate it to the nucleus to cleave defense related plant proteins. However, no other phytopathogen effectors have been shown to remove NLS from host proteins to disrupt their functions.

While the greatest amount of MED16 cleavage product was detected in the presence of virus and aphids (Fig. 3b), and in the presence of NIa-Pro and aphids (Fig 3d), the cleavage product was also detected in transgenic control plants infested with aphids alone (Fig 3d). In contrast to this, the MED16 cleavage product was not detected in the mock-inoculated plants with aphids using total protein extracts (Fig. 3b), however it was detected in this treatment when cytosolic protein fractions were used (Fig. 4c). This suggests that either some unknown aphid effector or an aphid-induced plant protease can also cleave MED16. The MED16 cleavage product was also detected in total protein extracts of EV control plants (Fig. 3d), however they were not detected in the cytosolic extracts (Fig. 4d). Potentially this suggests cleavage in these samples happened after protein extraction, or this may suggest there is a house-keeping role for MED16 cleavage in plants that is yet to be understood.

Plants, insects, and microbes have been co-evolving for millions of years. During this process, they have developed the ability to recognize each other as either beneficial partners or antagonists. As plants cannot move, they rely heavily on recognition of chemical signals from microbes and insects to differentiate between beneficial and harmful interactions and respond accordingly. One type of signal plants have evolved to recognize are the peptides created when pathogen and herbivore proteins are broken down by the host plant or by the colonizing organisms themselves (Marmiroli and Maestri, 2014; Albert, 2013). It was shown that cells of tomato and Arabidopsis plants can perceive nanomolar concentrations of only a 15 amino acid fragment of the bacterial flagellin protein (Mueller *et al*., 2012). Plants can also recognize Inceptin, a 11 amino acid cleaved peptide fragment of chloroplast ATPase, that was shown to induce defenses against caterpillars in cowpea (Schmelz *et al*., 2007). It is not known if MED16 cleavage products are recognized by host plants and used as a signal to upregulate defenses in adapted hosts. However, this work paves the way for future work on signal recognition and defense response in plant-virus-interactions, as well as on regulation of the MED16 protein.

## Acknowledgements

This work was supported by NSF grant # IOS-2026068 awarded to CLC. We would like to thank Dr. Zhang for sharing *med16* mutant seeds and the MED16-FLAG complemented Arabidopsis lines. We would like to thank Lior Lisk for assistance with experiment set-up, Ethan Eddy for technical support in the lab, and Chunling Yang for making the NIa-Pro protease mutant constructs and transgenic Arabidopsis plants.

---

**Figure S1:**
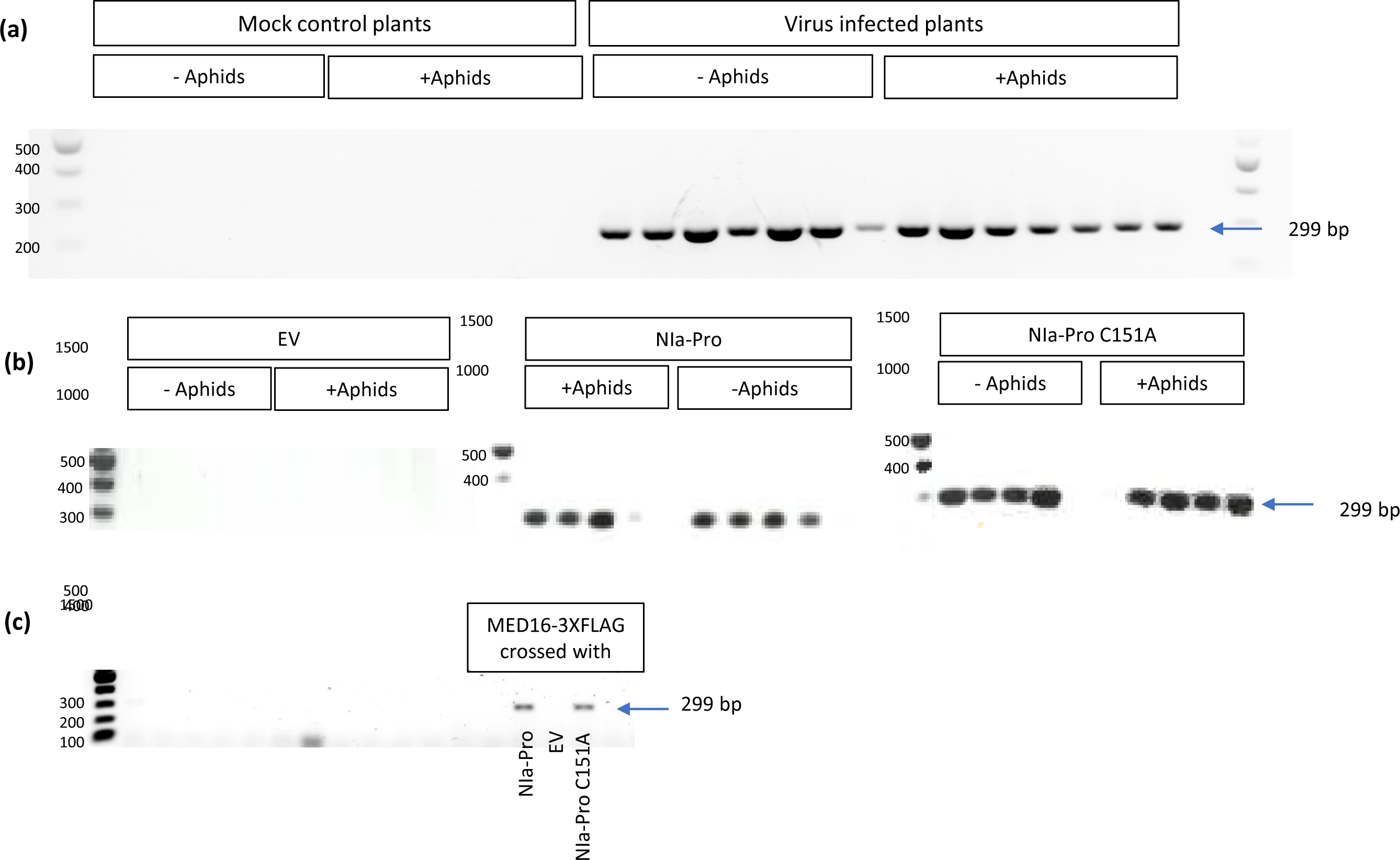
RT-PCR gel showing presence of NIa-Pro transcript in (a) Arabidopsis Col-0 plants that were rub-inoculated with TuMV-GFP, (b) Arabidopsis plants that were overexpressing the empty plasmid vector (EV), NIa-Pro or NIa-Pro C151 mutant, or (c) Arabidopsis mutants complemented with MEG16-3XFLAG and crossed with plants overexpressing the empty plasmid vector (EV), NIa-Pro, or NIa-Pro C151 mutant for the T2 generation (pool of N=6). The amplified product from C151A was sequenced to verify the C151A mutation in the NIa-Pro mutant over-expressor. The primers used amplify a 299 bp amplicon of NIa-Pro from TuMV at the 5’-end of the gene.

**Figure S2:**
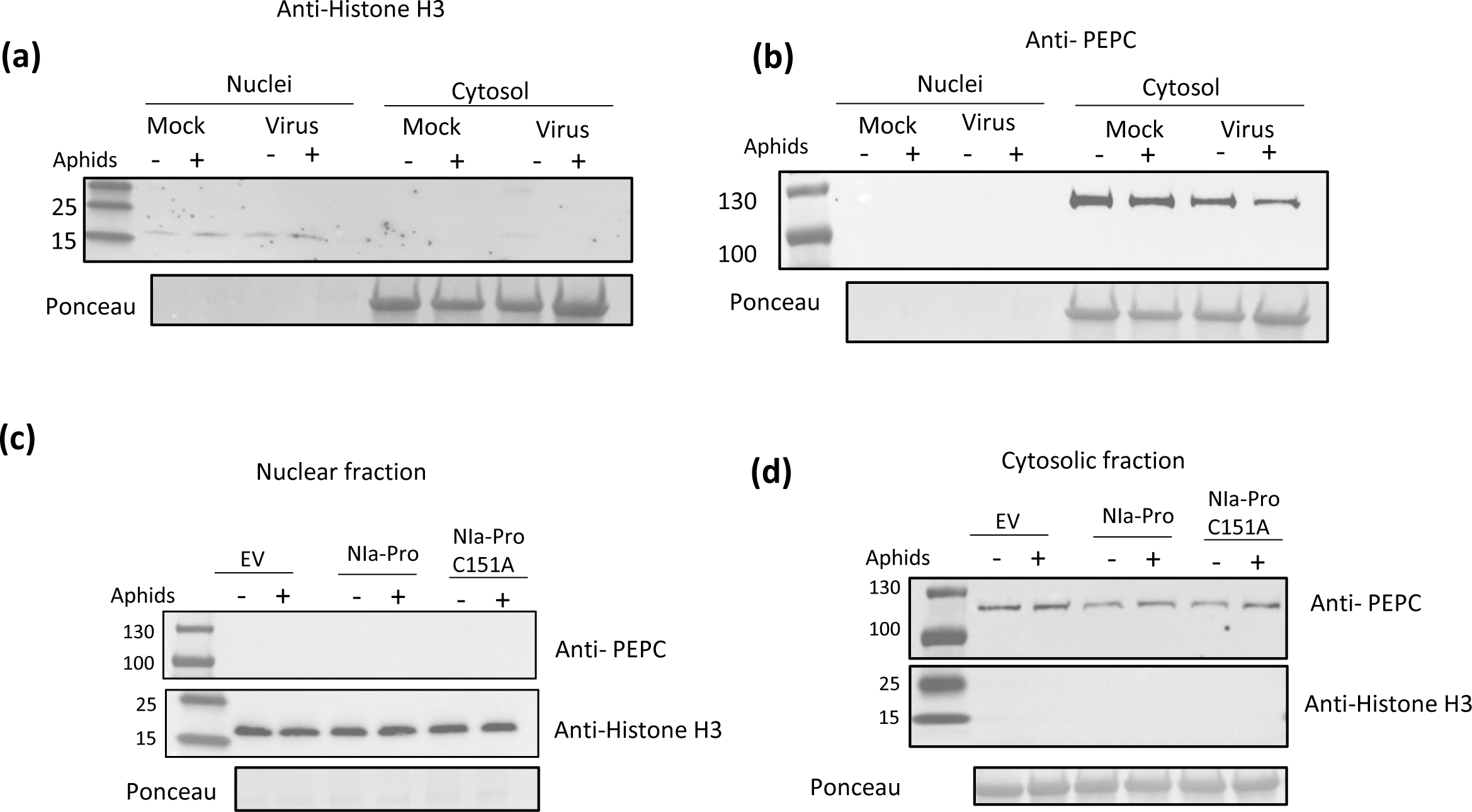
Proteins extracted from the cytosolic and nuclear fractions of Arabidopsis plants that were (a, b) mock or virus-infected or (c, d) expressing the empty expression vector (EV), NIa-Pro, or the protease mutant NIa-Pro C151A, and expressing MED16:3XFLAG. Pure organellar separations were verified by immunodetection of (a, c) the nucleus specific ∼17kD histone H3 protein histone H3 antibodies and (b, d) the cystosol specific ∼105kD phosphoenol pyruvate carboxylase or PEPC (b, d).

**Figure S3:**
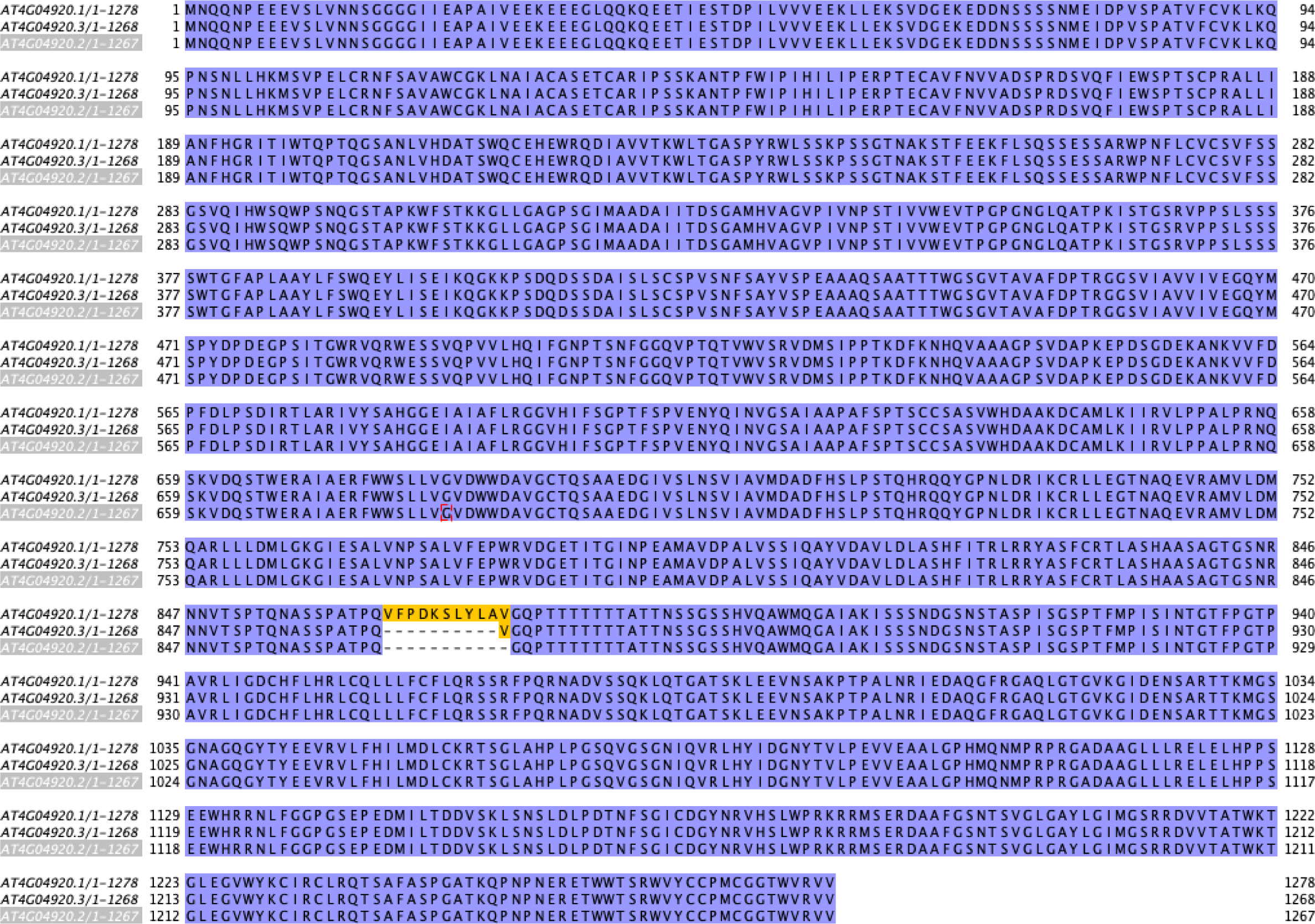
Sequence comparison of the three predicted isoforms of MED16 (AT4G04920)

**Figure S4:**
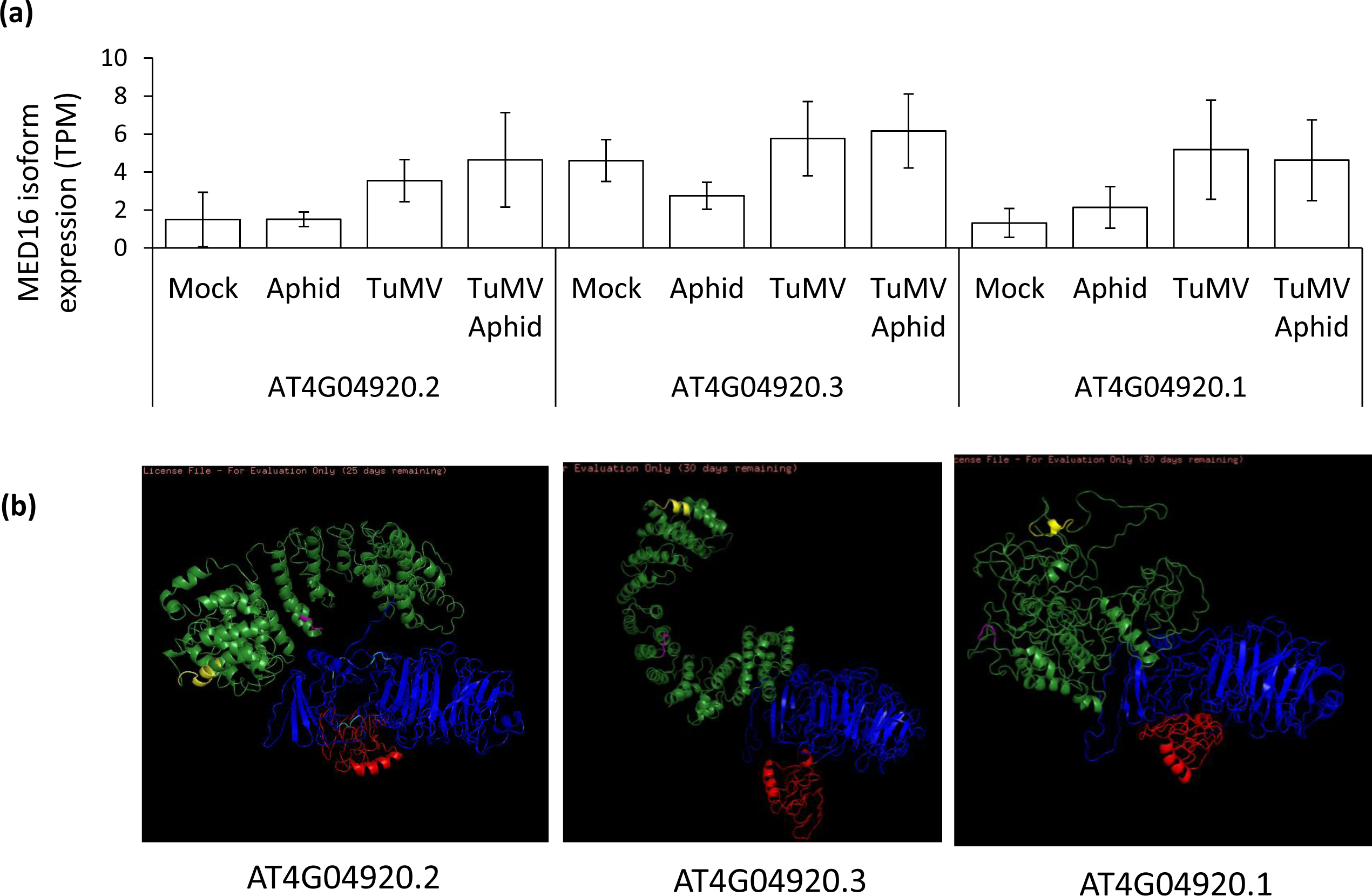
AT4G04920.3 is the most abundant isoform of MED16. (a) Isoform abundance of MED16 AT4G04920.1, AT4G04920.2, and AT4G04920.3 in Arabidopsis plants with and without TuMV infection and aphid feeding. (b) The predicted protein structures of the different MED16 isoforms (AT4G04920.1, AT4G04920.2, and AT4G04920.3) using I-TASSER.

**Table S1:**
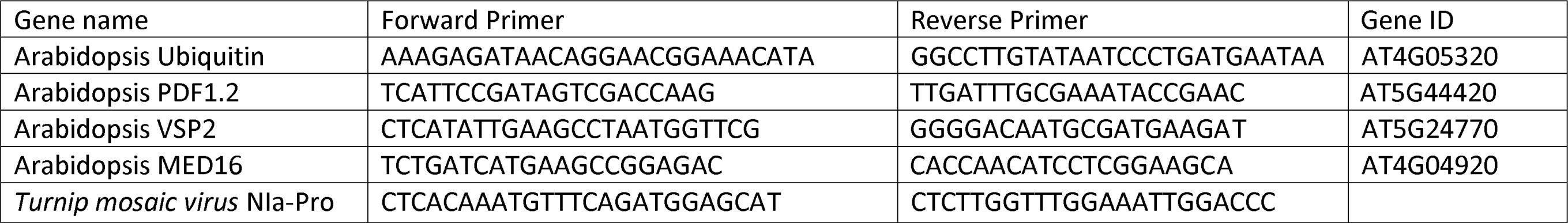
List or primers used:

